# Antibody Responses Against Coronaviruses in the Wuhan Population

**DOI:** 10.64898/2026.02.15.705976

**Authors:** Shixiong Zhou, Chunhai Liu, Weiyong Liu, Kun Cai, Ke Lan, Chengliang Zhu, Qian Wang, Lihong Liu

## Abstract

Prolonged circulation of SARS-CoV-2 may broaden population-level neutralizing antibody responses to diverse coronaviruses; however, the extent of this breadth remains unclear. To address this gap, we assessed serum neutralizing activity in 869 individuals from Wuhan, China, including 78 pre-pandemic samples collected in 2017 and 791 post-pandemic samples obtained in 2025. Specifically, using pseudovirus neutralization assays, sera were tested against a panel of coronaviruses, including circulating SARS-CoV-2 Omicron subvariants XFG and NB.1.8.1. Neutralization was measured at a fixed dilution for all samples and by determination of 50% inhibitory dilution (ID_50_) for selected sera. In parallel, antigenic cartography and genetic distance analyses were applied to relate functional antigenic relationships to spike protein divergence. Our results showed that, compared with pre-pandemic sera, post-pandemic sera exhibited enhanced neutralizing activity against multiple sarbecoviruses and the merbecovirus MjHKU4r-CoV-1, with mean inhibition increases ranging from 1.7% to 76.5% at a 1:20 dilution. Notably, the largest increases from 51.8% to 76.5% were observed against clade 1b sarbecoviruses. In contrast, neutralizing activity against human alphacoronaviruses and MERS-CoV showed no meaningful enhancement. Moreover, neutralizing titers in post-pandemic sera were strongly correlated across sarbecoviruses, particularly among clade 1b viruses. Antigenic mapping further revealed that several zoonotic sarbecoviruses were antigenically closer to ancestral SARS-CoV-2 than contemporary Omicron subvariants, despite substantially greater genetic divergence. Taken together, these findings demonstrate that cumulative SARS-CoV-2 exposure promotes broadly cross-neutralizing antibody responses within the sarbecovirus subgenus and provide a quantitative framework to inform the rational design of pan-sarbecovirus vaccines and antibody-based countermeasures for future coronavirus preparedness.

## Main text

Over the past two decades, the emergence of highly pathogenic coronaviruses, including SARS-CoV-1, MERS-CoV, and SARS-CoV-2, has highlighted their capacity to cross species barriers and sustain transmission in humans^1^. The continued global circulation of SARS-CoV-2 has resulted in extensive population exposure over time, providing prolonged immunological stimulation that may induce cross-neutralizing antibody responses against diverse coronaviruses^2^. Accordingly, characterizing humoral immune landscapes is essential for defining how cumulative viral encounters shape herd immunity and for guiding vaccine optimization against both existing and emerging coronavirus threats.

To address this question, we evaluated serum virus-neutralizing activity in 791 patients with influenza-like illness (ILI) recruited in Wuhan between May and October 2025 and compared these data with those from 78 individuals without known respiratory infections sampled in 2017, before the COVID-19 outbreak. All individuals in the post-pandemic cohort were screened for SARS-CoV-2 infection at the time of sampling by reverse-transcription quantitative polymerase chain reaction (RT-qPCR). Approximately 9% tested positive, a proportion consistent with the SARS-CoV-2 positivity among ILI cases reported by the Chinese Center for Disease Control and Prevention (China CDC) during the same timeframe^3^. Those in the post-pandemic cohort had received varying numbers of inactivated SARS-CoV-2 vaccine doses and had experienced prior SARS-CoV-2 infections, although precise exposure histories were not fully ascertainable, reflecting real-world conditions. Detailed clinical characteristics of both cohorts are summarized in **Table S1**.

We then quantified neutralizing antibody activity at a fixed serum dilution of 1:20 using a pseudovirus neutralization assay against a panel of diverse coronaviruses, including viruses reported to utilize human ACE2, DPP4, or APN as entry receptors^4^ (**Figure S1A**). The viral set comprised alphacoronaviruses HCoV-NL63 and HCoV-229E, merbecoviruses MERS-CoV and MjHKU4r-CoV-1, and multiple sarbecoviruses, with particular emphasis on the currently predominant SARS-CoV-2 Omicron subvariants, XFG globally and NB.1.8.1 in China (**Figure S1B**). A clade 4 sarbecovirus RmYN05 which uses bat Rmal ACE2 was also included to expand the diversity of viruses examined (**Figure S1A**). The cell lines for pseudovirus neutralization assays are summarized in **Figure S2A**. For Khosta-2, 293T cells expressing the natural host receptor, bat Raff ACE2, were used because of poor infectivity via human ACE2 (**Figures S2B and S2C**). On this basis, the receptor-usage framework enables a systematic evaluation of neutralizing antibody responses across genetically and functionally heterogeneous coronaviruses. Using this approach, we observed that post-pandemic sera exhibited enhanced neutralizing activity against sarbecoviruses clades 1a, 1b, 3, and 4, as well as the merbecovirus MjHKU4r-CoV-1, with increases in mean inhibition ranging from 1.7% to 76.5% (**Figure 1A**) at a 1:20 dilution. Notably, the most pronounced effects were observed for clade 1b sarbecoviruses, including the SARS-CoV-2 variants D614G, XFG, and NB.1.8.1, for which mean inhibition increased by 51.8% to 76.5%. In contrast, neutralizing activity against HCoV-229E, HCoV-NL63, and MERS-CoV showed no significant change. Unexpectedly, both the pre- and post-pandemic cohorts robustly neutralized Khosta-2, with mean inhibitions of 96.7% and 98.4%, respectively at a 1:20 serum dilution. To investigate the underlying mechanism, we analyzed 12 sera from each cohort. Sera from both groups efficiently bound to and neutralized Khosta-2, with higher activity observed in post-pandemic samples (**Figure S3A and S3B**). A chimeric pseudovirus neutralization assay further demonstrated that the neutralizing activity against Khosta-2 was primarily directed toward the S1 subunit (**Figure S3C**). In parallel, sequence alignment revealed >41% amino acid identity between the Khosta-2 spike protein and those of four seasonal human coronaviruses (**Figure S3D**). Both cohorts exhibited binding (**Figure S3E**) or neutralizing (**Figure S3F**) antibody responses to four seasonal coronaviruses, suggesting that prior seasonal coronavirus infections may contribute to the Khosta-2 neutralization observed in pre-pandemic sera.

**Figure 1.**
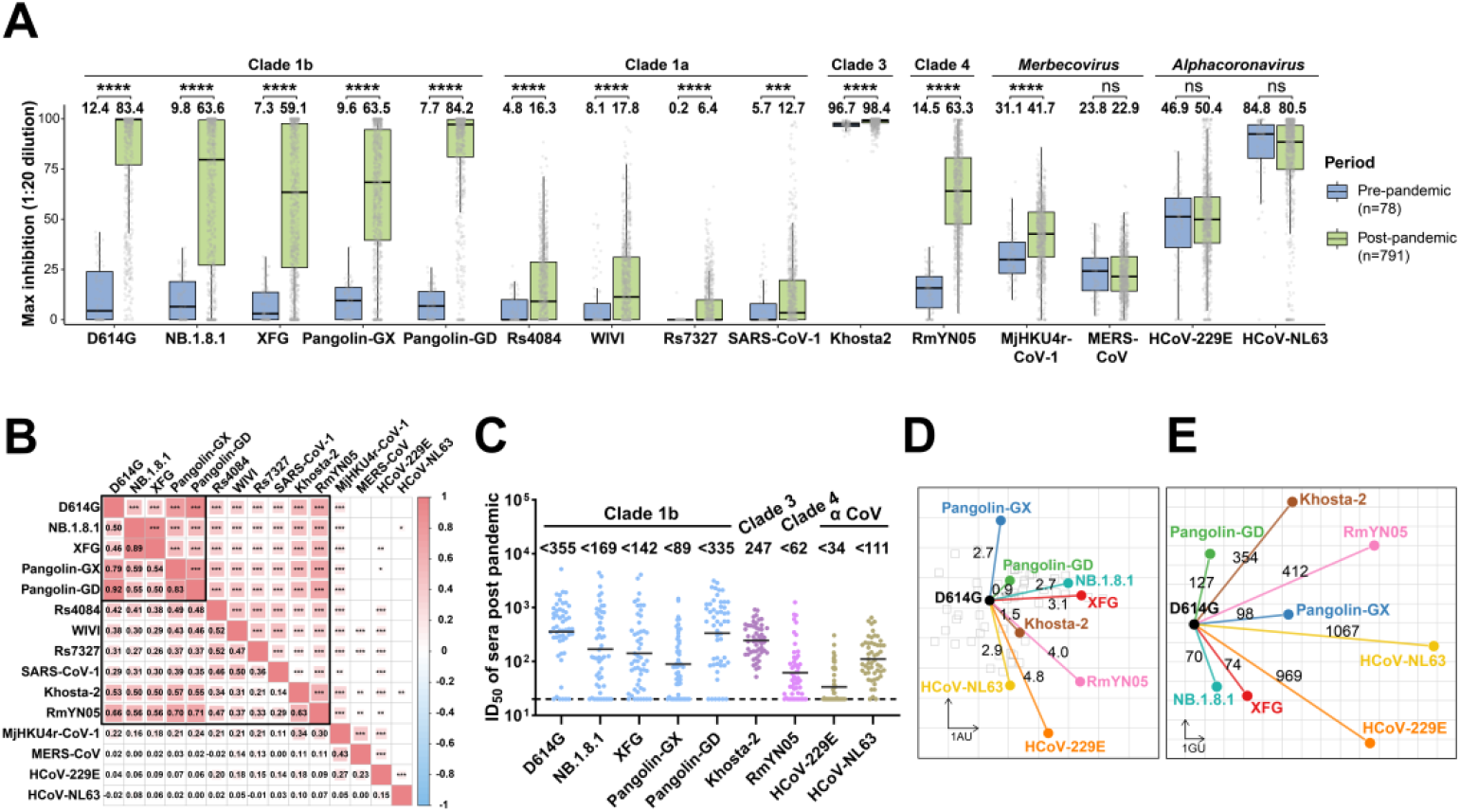
Neutralization profile of serum samples against coronaviruses in the Wuhan population. **A**. Neutralization of pseudotyped viruses in pre- and post-pandemic sera. Serum samples collected before (pre-pandemic, n=78) and after (post-pandemic, n=791) the pandemic were tested at a 1:20 dilution. Viruses are stratified by subgenus and clade: *Sarbecovirus* (clades 1b, 1a, 3, and 4), *Merbecovirus*, and *Alphacoronavirus*. Data are presented as box plots, where the horizontal line indicates the median and the box boundaries represent the interquartile range. Numerical values above the whiskers denote the mean inhibitions for the respective groups. Statistical significance was determined using the Wilcoxon rank-sum test, with *P* values adjusted by the Benjamini-Hochberg method (\*\*\**P* < 0.001; \*\*\*\**P* < 0.0001; ns, not significant). **B**. Heatmap of pairwise Spearman’s rank correlation coefficients (*r*_*s*_) for inhibitions against pseudotyped viruses. The analysis was based on post-pandemic serum samples in **A**. Correlation coefficients are displayed as numerical values in the lower triangle, while the upper triangle indicates statistical significance with asterisks (\**P* < 0.05, \*\**P* < 0.01, \*\*\**P* < 0.001). The size and color intensity of squares correspond to the *r*_*s*_ values, with colored squares representing significant correlations. **C**. Neutralization ID_50_ titers of 52 selected serum samples from the post-pandemic cohort. The dotted horizontal line marks the assay detection threshold. Geometric mean ID_50_s are denoted. **D**. An antigenic cartography was constructed using neutralization ID_50_s from panel **C**. D614G was used as the central reference, and antigenic distances were calculated as the mean divergence from each virus. One antigenic unit (AU) corresponds to an approximate twofold change in ID_50_ titer. Serum samples and viruses are represented by squares and dots, respectively. **E**. “Genetic map” constructed from pairwise spike amino acid edit distances between viral strains. D614G was used as the central reference. One genetic unit (GU) represents a log2-transformed amino acid difference.

Moreover, analysis of inhibitions at a 1:20 serum dilution revealed that serum neutralizing activities against sarbecoviruses were positively associated after pandemic, with the strongest correlations observed among clade 1b viruses (**Figure 1B**). Furthermore, neutralizing activity against a subset of viruses in this study declined with increasing age, as reflected by odds ratios of less than 1 (**Figure S4A**), underscoring the necessity for vaccine boosting to protect elderly susceptible populations. In parallel, sera from female participants exhibited elevated neutralizing activity against clade 1b, clade 3, and clade 4 sarbecoviruses compared with sera from male participants (**Figure S4B**). However, the underlying explanation for this observation remains unclear.

To extend our analysis of neutralizing titers, we selected 52 post-pandemic sera that exhibited pronounced neutralizing activity against ancestral SARS-CoV-2 D614G at a 1:20 dilution for determination of the 50% inhibitory dilution (ID_50_). The selected samples showed similar distributions of age, sex, and COVID-19 positivity compared with the overall post-pandemic cohort (**Table S2**). ID_50_ values against clade 1a sarbecoviruses and merbecoviruses were not determined because most sera did not achieve 50% inhibition at this dilution. Overall, geometric mean ID_50_ values for sarbecoviruses tested ranged from approximately 62 to 355, whereas those for HCoV-229E and HCoV-NL63 were lower, at approximately 34 and 111, respectively (**Figure 1C**). Consistent with this pattern, neutralizing titers against SARS-CoV-2 D614G were positively correlated with titers against other sarbecoviruses, but not with those against the non-sarbecovirus coronaviruses tested (**Figure S5**). An antigenic map was then generated on the basis of neutralization ID_50_ values, revealing that the sarbecoviruses Pangolin-GD, Pangolin-GX, and Khosta-2 were antigenically closer to D614G than the SARS-CoV-2 variants XFG and NB.1.8.1 (**Figure 1D**). In contrast, amino acid-based genetic distance analysis showed the opposite pattern, with these viruses exhibiting substantial genetic divergence from D614G (**Figure 1E**). Together, our results align with previous work showing that antigenic relatedness does not strictly track genetic divergence, underscoring the functional dominance of conserved neutralizing epitopes^5^. Collectively, we performed large-scale serum neutralization assays and constructed an antigenic map, enabling direct quantitative comparison of antigenic similarity between circulating SARS-CoV-2 strains and genetically divergent coronaviruses. Our findings indicate that cumulative SARS-CoV-2 exposure drives the development of broadly cross-neutralizing antibody responses against distinct coronaviruses, particularly within the sarbecovirus subgenus, implicating shared cross-reactive antigenic epitopes across these viruses. In summary, these results establish a practical framework for prioritizing viral targets in vaccine design and for leveraging existing population immunity to strengthen preparedness against ongoing SARS-CoV-2 evolution and future coronavirus emergence.

## Supplementary Methods

### Clinical cohorts

A total of 78 pre-pandemic serum samples were collected in Wuhan, China, between May and July 2017 from individuals without known respiratory infections, and 791 post-pandemic serum samples were collected in the same city between May and October 2025 from individuals with influenza-like illness (ILI). Immunocompromised patients were excluded from both cohorts. All post-pandemic samples were screened by reverse transcription quantitative polymerase chain reaction (RT-qPCR) to determine acute SARS-CoV-2 infection status at the time of sampling, and all participants in both cohorts provided written informed consent under protocols approved by the Hubei Provincial Center for Disease Control and Prevention (Approval No.: HBCDC-AF/SC-08/02.0) and Tongji Hospital (Approval No.: TJ-IRB20191225). Clinical information of these two cohorts are summarized in **Table S1**, and detailed characteristics of the 52 selected post-pandemic samples shown in **Figure 1C** are provided in **Table S2**.

### Phylogenetic analysis

Full-length spike glycoprotein sequences of the *Betacoronavirus* subgenus were retrieved from the GenBank and GISAID databases. The sequences were aligned using MAFFT (version 7.525)^1^. A maximum-likelihood phylogenetic tree was inferred using IQ-TREE software (version 3.0.1)^2^. The best-fit substitution model, WAG+F+I+R5, was selected by ModelFinder based on the Bayesian Information Criterion (BIC)^3^. Branch support was assessed using 1,000 ultrafast bootstrap replicates. The final tree was rooted using *Alphacoronavirus* sequences and subsequently visualized and annotated using tvBOT (version 2.6.1)^4^. The phylogenetic tree based on amino acid sequences of coronavirus spike proteins in this study is presented in **Figure S1**.

### Cell lines

HEK-293T cells (CRL-3216) were obtained from the American Type Culture Collection (ATCC) and used for pseudovirus production. For pseudovirus neutralization assays, the following target cell lines were used: Caco2, 293T-hACE2 (human ACE2), Vero E6-TMPRSS2-T2A-ACE2, 293T-RaffACE2, and 293T-RmalACE2; detailed information for all cell lines is provided in **Figure S2A**. Notably, the 293T-RaffACE2 and 293T-RmalACE2 cell lines were generated by lentiviral transduction, followed by selection with 1 μg/mL puromycin, and their ACE2 expression was confirmed by Western blotting (**Figure S2B**). All cell lines were cultured in Dulbecco’s modified Eagle medium (DMEM) supplemented with 10% fetal bovine serum (FBS) and 1% penicillin-streptomycin at 37 °C in a humidified atmosphere containing 5% CO2.

### Plasmids

Coronavirus spike-expressing plasmids (**Figure S2A**) were used for pseudovirus packaging. They were obtained from Yunlong Cao (Peking University, Beijing, China), Zheng-Li Shi (Guangzhou National Laboratory, Guangzhou, China), Qihui Wang and George F. Gao (Institute of Microbiology, Chinese Academy of Sciences, Beijing, China), or synthesized by BioSune Biotechnology (Shanghai, China), and cloned into the pCMV3 eukaryotic expression vector (Sino Biological Inc., Beijing, China). In addition, the Raff-ACE2-3×Flag and Rmal-ACE2-3×Flag plasmids, used to generate ACE2 stably expressing cell lines, were kindly provided by Dr. Huan Yan (Wuhan University). All constructs were confirmed by Sanger sequencing. To enhance pseudovirus infectivity, the C termini of multiple spike proteins were truncated: 18 amino acids were removed from D614G, XFG, NB.1.8.1, and Khosta-2; 17 amino acids from Pangolin-GX, Pangolin-GD, Rs4084, WIV1, Rs7327, SARS-CoV-1, and RmYN05; and 14 amino acids from HCoV-229E and HCoV-NL63. In contrast, the spike proteins of MjHKU4r-CoV-1 and MERS-CoV were used in their full-length forms.

### Pseudotyped virus

Pseudotyped viruses were generated as previously described^5^. Briefly, HEK-293T cells were seeded in 150 cm^2^ dishes one day before transfection. The following day, 30 μg of plasmids encoding coronavirus spike proteins were transfected using PEI-MAX (1 mg/mL). 24 hours post-transfection, cells were infected with VSV-G-pseudotyped ΔG-luciferase virus at a multiplicity of infection (MOI) of approximately 3-5 (G*ΔG-luciferase; Kerafast). After a 2-hour incubation, the inoculum was removed, and cells were washed three times with DMEM supplemented with 2% FBS and 1% penicillin-streptomycin. Subsequently, 20 mL of fresh medium was added. Pseudovirus-containing supernatants were harvested 24 hours later and clarified by centrifugation. To eliminate contaminating VSV-G pseudovirus, clarified supernatants were incubated with 20% I1 hybridoma supernatant (ATCC CRL-2700) before aliquoting and storage at -80 °C until use.

To investigate the potential epitopes recognized by neutralizing antibodies in sera against Khosta-2, two chimeric pseudoviruses were generated. Specifically, the SARS1-S1&Khosta2-S2 chimeric spike was constructed by fusing the SARS-CoV-1 S1 subunit (Met1-Arg667) to the Khosta-2 S2 subunit (Ala667-Lys1235). Conversely, the Khosta2-S1&SARS1-S2 chimeric spike was generated by fusing the Khosta-2 S1 subunit (Met1-Arg666) to the SARS-CoV-1 S2 subunit (Ser668-Lys1237). Pseudotyped viruses were then generated as described above.

### Pseudovirus neutralization assay

Pseudovirus neutralization assay was performed as previously descripted^6^. To ensure consistent viral input, all pseudoviruses were titrated before each round of neutralization assays. Serum samples were heat-inactivated at 56 °C for 30 min before the assay. For inhibition experiments, each serum sample was 20-fold diluted in culture medium. For determination of the half-maximal inhibitory dilution (ID_50_), sera were subjected to a series of four-fold serial dilutions starting from 1:20. Subsequently, 50 μL of each serum dilution was mixed with pseudovirus and incubated at 37 °C for 1 h. Next, 100 μL of target cell suspension was added to each well, and plates were incubated overnight at 37 °C in a humidified atmosphere containing 5% CO2. Luciferase activity was measured using a multifunctional microplate reader (Thermo Fisher Scientific).

### ELISA

The S2P-stabilized spike trimer of Khosta-2 was generated using a method similar to that previously described for construction of the SARS-CoV-2 D614G spike trimer^7^. The HCoV-OC43 and HCoV-NL63 spike proteins were kindly provided by Dr. Yan Gao (ShanghaiTech University)^8^. To determine serum binding titers to Khosta-2, HCoV-OC43, and HCoV-NL63 spike proteins, 96-well ELISA plates were coated overnight at 4°C with 50 ng per well of the respective S2P trimer proteins. Plates were then blocked with 300 µL of blocking buffer (10% FBS in PBST (0.1% Tween 20 in PBS)) at 37°C for 1 hour. Serum samples were serially diluted fourfold in dilution buffer (10% FBS in PBST), starting at 1:100, added to the plates, and incubated at 37°C for 1 hour. After washing with PBST, HRP-conjugated goat anti-mouse IgG (H+L) antibody (ABclonal) was added and incubated at 37°C for 1 hour. Plates were washed between each step. TMB substrate (in-house) was then added and incubated for 6 minutes, and the reaction was stopped with 2 M sulfuric acid. Absorbance was measured at 450 nm.

### Sequence identity analysis

Pairwise amino acid sequence identities were evaluated for the full-length spike protein, as well as the S1 and S2 subunits. Global alignments were performed using the Needleman-Wunsch^9^ algorithm implemented in the Biopython^10^ suite. Percent identity was calculated as the proportion of identical residues relative to the total length of the alignment.

### Antigenic cartography

Antigenic distances between sera and coronaviruses were inferred by integrating ID_50_ values from individual serum samples using a previously described antigenic cartography approach^11^. Antigenic maps were visualized with the Racmacs package (version 1.2.9; https://acorg.github.io/Racmacs/) implemented in the R environment (version 4.5.1). Map optimization was performed over 2,000 iterations, with the “minimum column basis” parameter set to “none.” Antigenic distances were calculated using the mapDistances function. D614G was designated as the central reference, and map orientation was refined by manual adjustment of seeds. One antigenic unit represents an approximately twofold change in ID_50_ titer.

### Genetic distance analysis

Genetic distances were calculated based on pairwise amino acid alignments using the Needleman-Wunsch algorithm^9^. Raw distances were log2-transformed and visualized via multidimensional scaling (MDS) using custom-written R scripts. The map layout was optimized using the L-BFGS-B algorithm, with D614G positioned as the center. One genetic unit corresponds to a 2-fold difference in amino acid substitutions.

### Quantification and Statistical analysis

Serum neutralization 50% inhibitory dilution (ID_50_) values were calculated by fitting the data to a nonlinear five-parameter dose-response model using GraphPad Prism software (version 10.1.2). Differences in neutralization activity between pre-pandemic and post-pandemic samples were assessed using the Wilcoxon rank-sum test, with P values adjusted for multiple comparisons using the Benjamini-Hochberg (BH) method. Correlations between serum inhibition at a 20-fold dilution against different viruses were evaluated using Spearman’s rank correlation. Logistic regression was used to calculate odds ratios (ORs) and to fit regression lines. Significance levels were annotated as follows: ns, not significant; \**P* < 0.05; \*\**P* < 0.01; and \*\*\**P* < 0.001, \*\*\*\**P* <0.0001.

## Acknowledgments

This study was supported by the National Natural Science Foundation of China (grant nos. 92569102, awarded to L.L., and 82372223, awarded to K.C.). Additional resources were provided by the Taikang Center for Life and Medical Sciences, Wuhan University, Wuhan, China, and by the State Key Laboratory of Virology and Biosafety, Wuhan University, Wuhan, China, for L.L. and Q.W. This research also benefited from the Hubei Provincial Public Health Outstanding Young Talents Project and the Chu Tian Excellence Program (Medical and Health). We further express our appreciation to all contributors to the Global Initiative on Sharing All Influenza Data (GISAID). The sources of support had no role in the study design, data collection, analysis, decision to publish, or preparation of the manuscript. Finally, we sincerely thank David D. Ho, Ian A. Mellis, and Yicheng Guo of Columbia University, USA, for their valuable insights and constructive feedback on the manuscript.

## Author Contributions

Q.W. and L.L. conceived and supervised the project. Q.W., L.L., and S.Z. conducted the experiments. K.C., W.L., and C.Z. collected clinical samples and performed reverse-transcription quantitative polymerase chain reaction (RT–qPCR) assays to determine acute SARS-CoV-2 infection status. S.Z. managed the project and collected serum samples. C.L. performed the bioinformatic analyses. Q.W., L.L., S.Z., C.L., and K.L. analyzed the data and wrote the manuscript.

## Declaration of Interests

All authors declare no competing interests.

**Figure S1.**
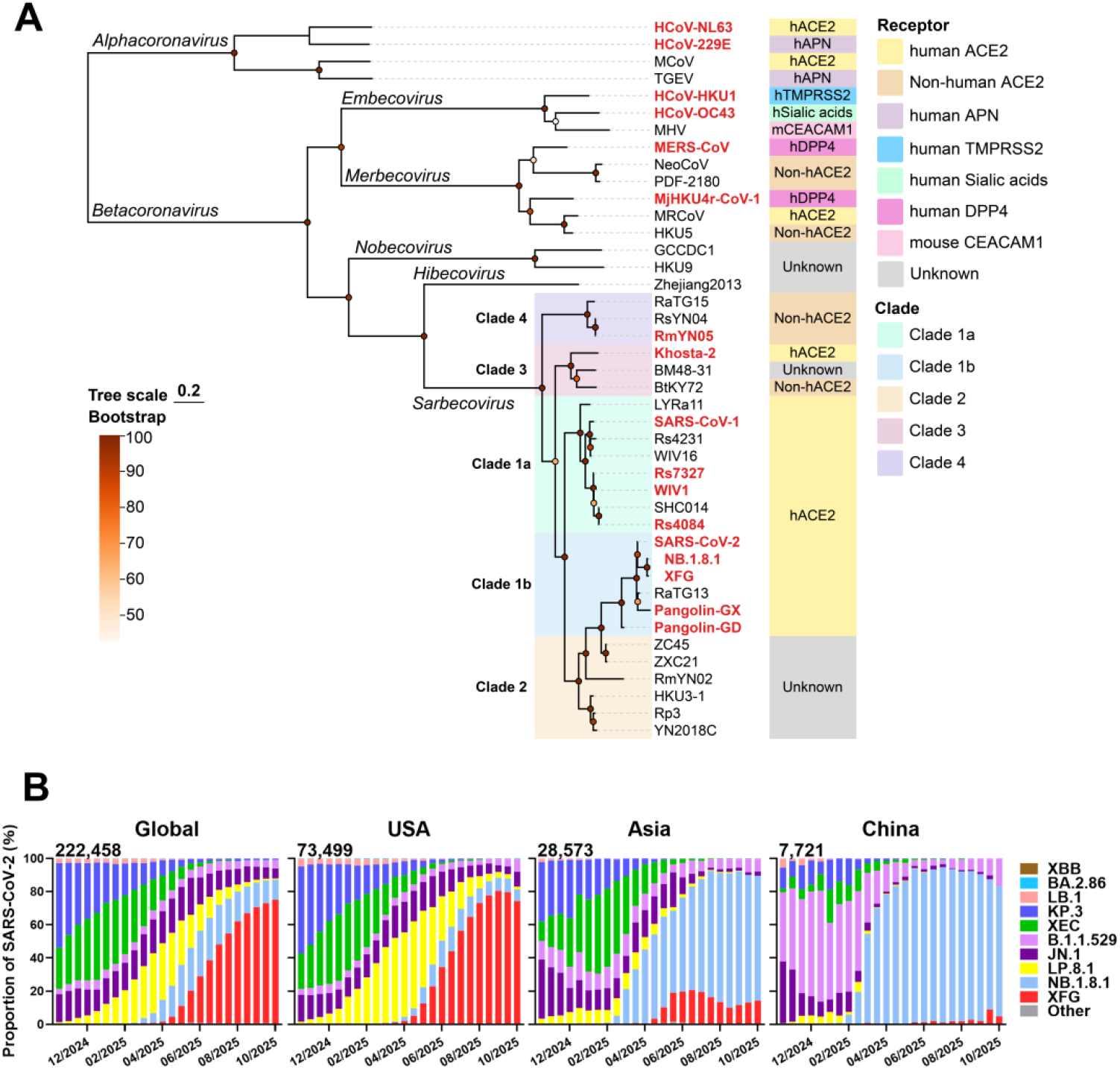
Evolutionary relationships of coronavirus spike proteins and prevalence of SARS-CoV-2 Omicron subvariants. A. Phylogenetic tree based on amino acid sequences of coronavirus spike proteins. Viruses included in the serum neutralization assays in this study are highlighted in red. B. Frequencies of SARS-CoV-2 subvariants during the indicated period. Raw data were obtained from the Global Initiative on Sharing All Influenza Data (GISAID). Values in the upper left corner of each panel denote the cumulative number of SARS-CoV-2 sequences deposited.

**Figure S2.**
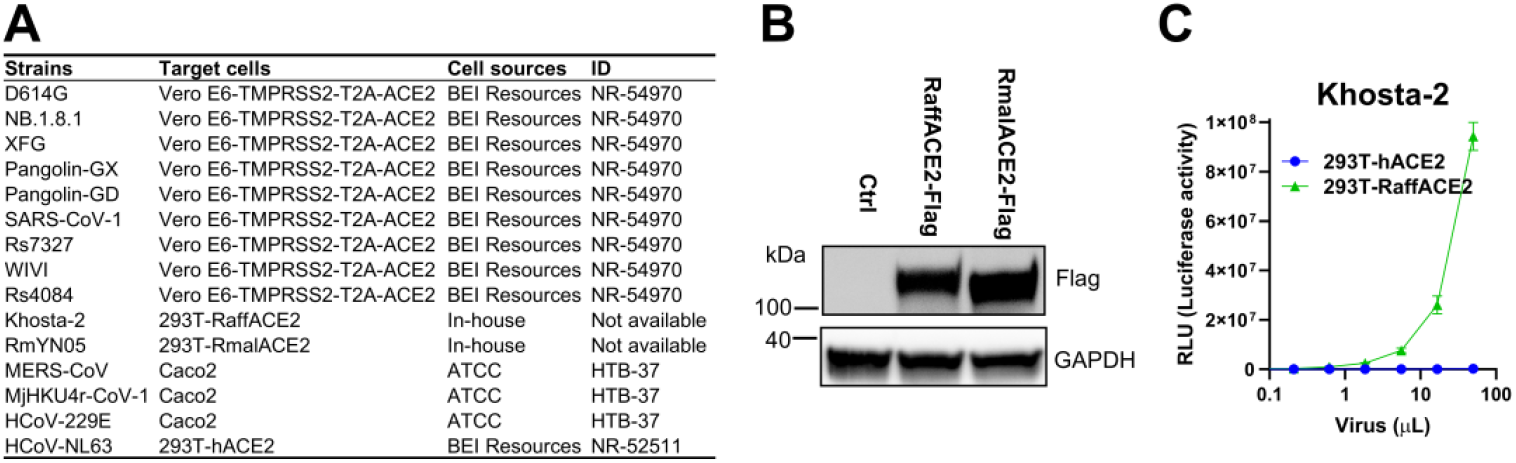
Summary of cell lines used for pseudovirus neutralization assays in this study. A. Virus-susceptible cell lines used for the indicated pseudoviruses. ATCC, American Type Culture Collection; BEI Resources, Biodefense and Emerging Infections Research Resources Repository. B. Immunoblot analysis of ACE2 expression on 293T-RaffACE2 and 293T-RmalACE2 cells. Wild-type 293T cells were used as a control (Ctrl). GAPDH served as an internal loading control. C. Infectivity of Khosta-2 pseudotyped virus in 293T-hACE2 and 293T-RaffACE2 cells.

**Figure S3.**
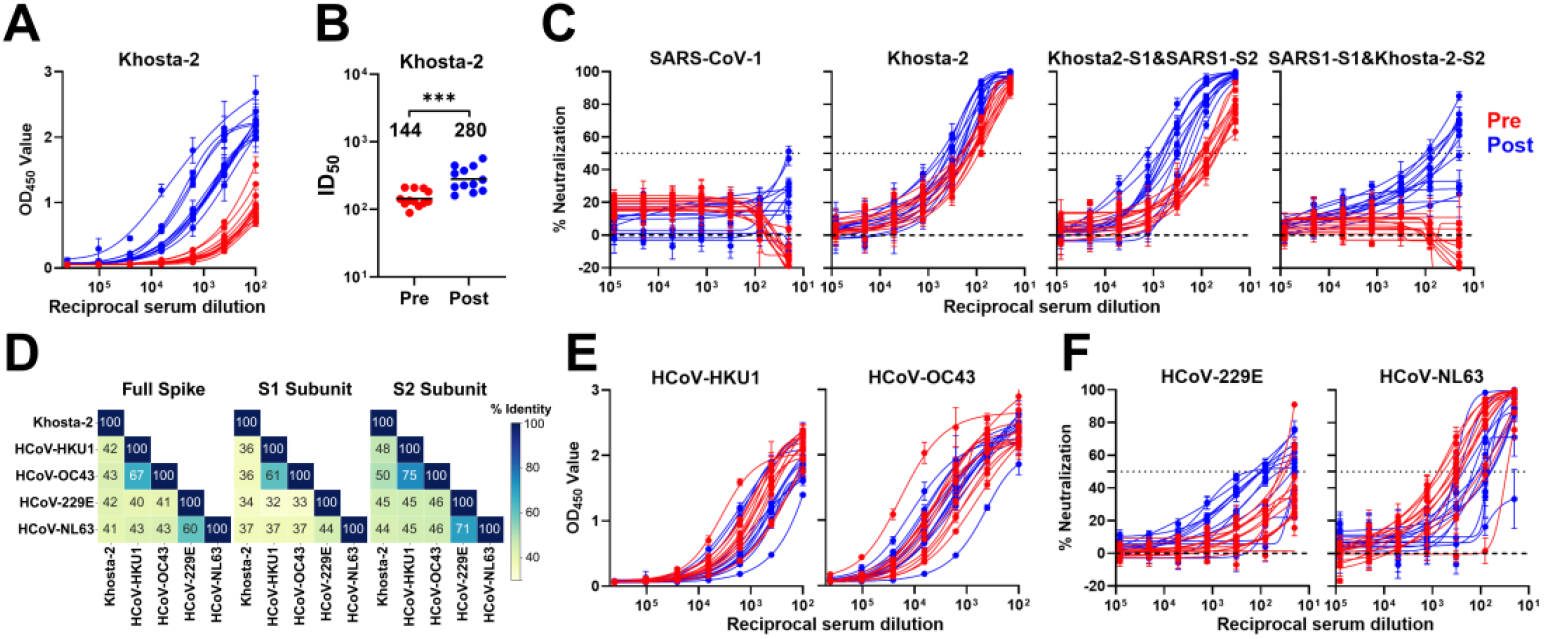
Binding and neutralization profiles of khosta-2 compared with other human coronaviruses. A. Binding curves of the Khosta-2 spike protein to serum samples from the pre- and post-pandemic cohorts. B. Neutralization ID_50_ values against Khosta-2 in serum samples from the pre- and post-pandemic cohorts. Statistical analysis was performed using an unpaired t-test. \*\*\**P* < 0.001. C. Neutralization curves of pseudotyped SARS-CoV-1, Khosta-2, and the indicated chimeric viruses by serum samples from the pre- and post-pandemic cohorts. SARS-CoV-1 was selected to generate chimeric viruses with Khosta-2 because it was resistant to neutralization by sera from both cohorts. D. Pairwise amino acid sequence identity of coronavirus spike proteins. Heatmaps display the percentage of amino acid identity between the spike proteins of Khosta-2, HCoV-HKU1, HCoV-OC43, HCoV-229E, and HCoV-NL63. Comparisons were performed for the full-length spike protein (left), the S1 subunit (middle), and the S2 subunit (right). The color scale indicates the percent identity, ranging from yellow (lower identity) to dark blue (higher identity). E. Binding curves of the HCoV-HKU1 and HCoV-OC43 spike proteins to serum samples from the pre- and post-pandemic cohorts. F. Neutralization curves of pseudotyped HCoV-229E and HCoV-NL63 by serum samples from the pre- and post-pandemic cohorts. Pre- and post-pandemic samples are shown in red and blue, respectively. Twelve samples were tested in each cohort.

**Figure S4.**
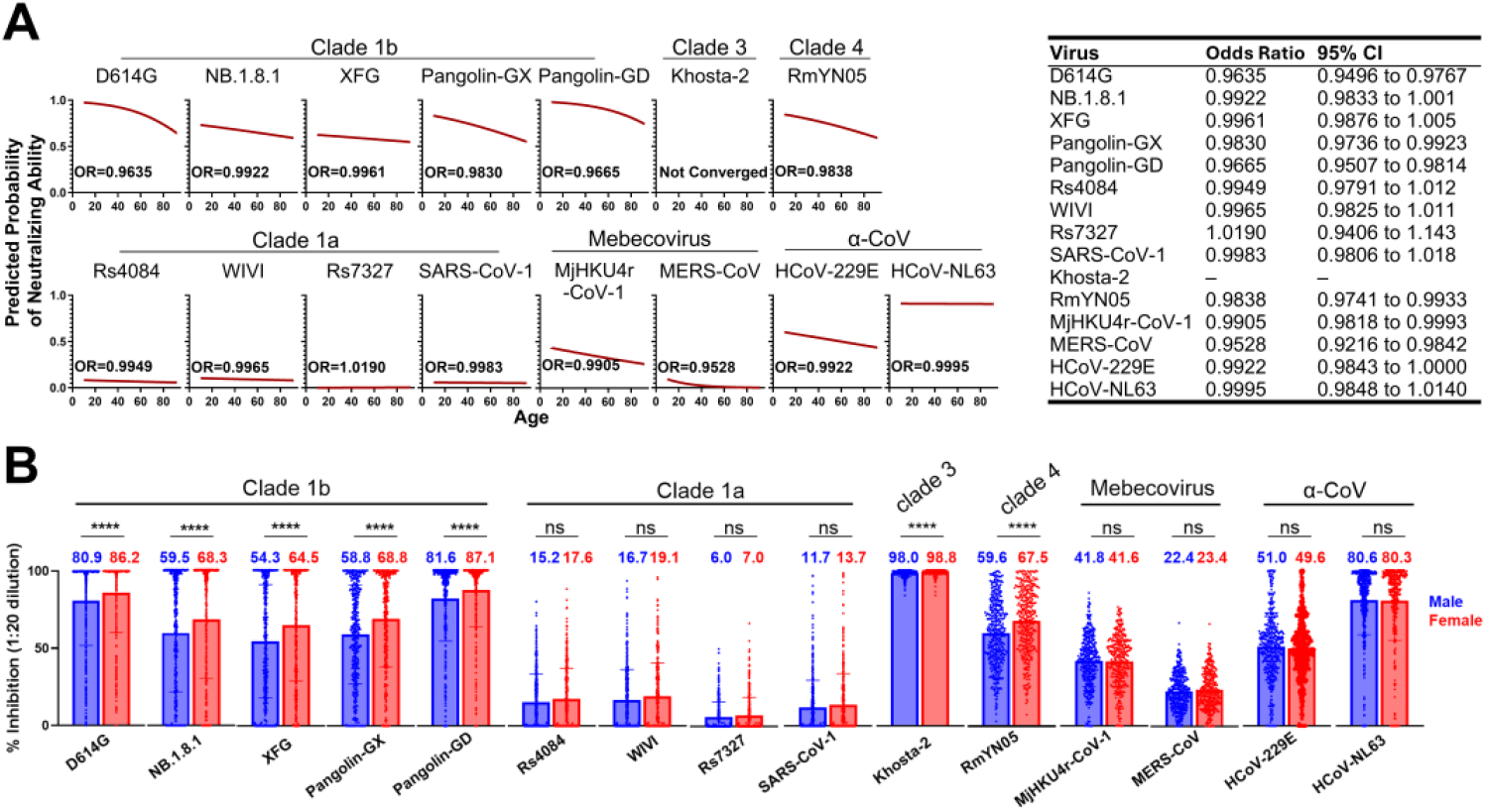
Effects of age and sex on the cross-neutralizing activity of sera against coronaviruses. A. Association between participant age and serum inhibition (at a 1:20 dilution) of the indicated pseudotyped viruses. Odds ratios (ORs) and 95% confidence intervals (CIs) are shown. -, data not available. B. Comparison of serum inhibition between sexes against the indicated pseudotyped viruses. Statistical significance was assessed using the Mann-Whitney U test (\*\*\*\**P* < 0.0001; ns, not significant). Values above the data points indicate mean inhibitions of sera at a 1:20 dilution. Inhibitions by sera from male participants are shown in blue, and that from female participants is shown in red.

**Figure S5.**
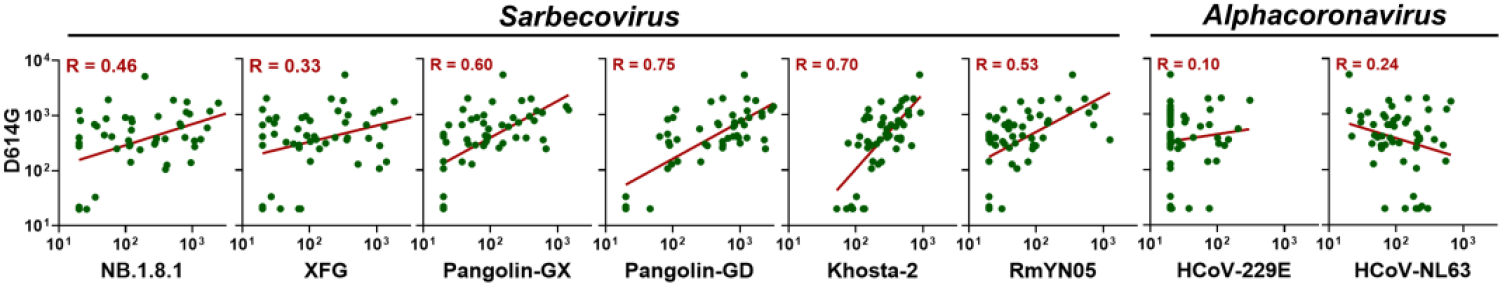
Correlation between serum neutralizing ID_50_ titers against SARS-CoV-2 D614G and other coronaviruses. Pearson correlation R values are shown.

**Table S1.** Summary of clinical cohorts in this study.

|  | **Pre-Pandemic**<br>**(n=78)** | **Post-Pandemic**<br>**(n=791)** |
| --- | --- | --- |
| **Age** |  |  |
| <46 | 39 (50.0%) | 140 (17.7) |
| 46-70 | 37 (47.5%) | 396 (50.1%) |
| >70 | 2 (2.5%) | 255 (32.2%) |
| **Sex** |  |  |
| Male | 43 (55.1%) | 423 (53.5%) |
| Female | 35 (44.9%) | 368 (46.5%) |
| **COVID-19 tested by RT-qPCR** |  |  |
| Positive | 0 (0%) | 71 (9%) |
| Negative | 78 (100%) | 720 (91%) |

RT-qPCR, reverse transcription quantitative polymerase chain reaction. n, sample size.

**Table S2.** Clinical characteristics of 52 selected post-pandemic samples for neutralization ID_50_ determination.

| Sample ID | Gender | Age | COVID-19 (RT-qPCR) | Race |
| --- | --- | --- | --- | --- |
| G3-21 | Male | 44 | Negative | Asian |
| G3-22 | Male | 51 | Negative | Asian |
| G3-23 | Male | 45 | Negative | Asian |
| G3-24 | Male | 40 | Negative | Asian |
| G3-25 | Male | 47 | Negative | Asian |
| G3-26 | Male | 45 | Negative | Asian |
| G3-27 | Male | 48 | Negative | Asian |
| G3-28 | Male | 43 | Negative | Asian |
| G3-29 | Male | 52 | Negative | Asian |
| G3-30 | Male | 48 | Positive | Asian |
| G3-31 | Female | 48 | Negative | Asian |
| G3-32 | Female | 43 | Negative | Asian |
| G3-33 | Female | 43 | Negative | Asian |
| G3-34 | Female | 55 | Negative | Asian |
| G3-35 | Female | 44 | Negative | Asian |
| G3-36 | Female | 40 | Negative | Asian |
| G3-37 | Female | 49 | Negative | Asian |
| G3-38 | Female | 53 | Negative | Asian |
| G3-39 | Female | 41 | Negative | Asian |
| G3-40 | Female | 48 | Positive | Asian |
| G3-41 | Male | 75 | Positive | Asian |
| G3-42 | Male | 54 | Negative | Asian |
| G3-43 | Male | 71 | Negative | Asian |
| G3-44 | Male | 74 | Negative | Asian |
| G3-45 | Male | 71 | Negative | Asian |
| G3-46 | Male | 61 | Negative | Asian |
| G3-47 | Male | 41 | Negative | Asian |
| G3-48 | Male | 40 | Negative | Asian |
| G3-49 | Male | 32 | Negative | Asian |
| G3-50 | Male | 55 | Negative | Asian |
| G3-51 | Male | 85 | Positive | Asian |
| G3-52 | Male | 79 | Negative | Asian |
| G3-53 | Male | 77 | Negative | Asian |
| G3-54 | Male | 40 | Negative | Asian |
| G3-55 | Female | 72 | Negative | Asian |
| G3-56 | Female | 61 | Negative | Asian |
| G3-57 | Female | 42 | Negative | Asian |
| G3-58 | Female | 71 | Negative | Asian |
| G3-59 | Female | 36 | Negative | Asian |
| G3-60 | Female | 65 | Negative | Asian |
| G3-61 | Female | 32 | Negative | Asian |
| G3-62 | Female | 46 | Negative | Asian |
| G3-63 | Female | 50 | Negative | Asian |
| G3-64 | Female | 50 | Negative | Asian |
| G3-65 | Female | 80 | Negative | Asian |
| G3-66 | Female | 51 | Negative | Asian |
| G3-67 | Female | 69 | Negative | Asian |
| G3-68 | Female | 56 | Negative | Asian |
| G3-69 | Female | 60 | Negative | Asian |
| G3-70 | Female | 74 | Negative | Asian |
| G3-71 | Female | 46 | Negative | Asian |
| G3-72 | Female | 77 | Positive | Asian |

